# Protozoan predation enhances stress resistance and antibiotic tolerance in the opportunistic pathogen *Burkholderia cenocepacia* by triggering the SOS response

**DOI:** 10.1101/2023.08.24.551519

**Authors:** Álvaro Morón, Iván Belinchón, Alaa E. Tarhouchi, Juan M. Valenzuela, Patricia de Francisco, Ana Martín-González, Francisco Amaro

## Abstract

Bacterivorous protists are thought to serve as training grounds for bacterial pathogens by subjecting them to the same hostile conditions that they will encounter in the human host. Bacteria that survive intracellular digestion exhibit enhanced virulence and stress resistance after successful passage through protozoa but the underlying mechanisms remain to be clarified. Here we show that the opportunistic pathogen *Burkholderia cenocepacia* survives phagocytosis by ciliates found in domestic and hospital sink drains, and viable bacteria are expelled packaged in respirable membrane vesicles with enhanced resistance to oxidative stress, desiccation and antibiotics, thereby contributing to bacterial dissemination in the environment. By using diverse methodological approaches, we demonstrate that reactive oxygen species generated within the protozoan phagosome promote the formation of persisters tolerant to ciprofloxacin by activating the bacterial SOS response. Besides, we show that genes encoding antioxidant enzymes are upregulated during passage through ciliates increasing bacterial resistance to oxidative radicals. We prove that suppression of the SOS response impairs bacterial intracellular survival and persister formation within protists. This study highlights the significance of protozoan food vacuoles as niches that foster bacterial adaptation in natural and built environments and suggests that persister switch within phagosomes may be a widespread phenomenon in bacteria surviving intracellular digestion.

## Introduction

Protozoan predation represents a major selection pressure that bacteria face in both natural and anthropogenic environments [1, 2]. Bacterivorous protozoa (traditionally termed as protozoa) such as ciliates, flagellates and amoebae, tightly regulate bacterial populations, but may also act as “Trojan horses” serving as reservoir where pathogenic bacteria can survive and even replicate [3, 4]. Macrophages and protozoa kill bacteria by taking them up into phagosomes that become acidified and filled with ROS and hydrolytic enzymes. However, some bacterial species can evade intracellular digestion by employing different mechanisms [5], subsequently infecting their predators [6, 7], or are eventually released packaged in vesicles expelled by protists which are often named fecal pellets or expelled food vacuoles (EFVs) [8–11]. Interestingly, studies have shown that pathogenic bacteria released from protozoan food vacuoles exhibit enhanced resistance to harsh conditions, antibiotics and even increased virulence [11–13], suggesting that EFVs may act as vehicles for pathogen transmission. In fact, the size of EFVs (2-6 μm in diameter) falls within the range of respirable particles [10], and thus may reach the lung alveoli leading to infection, although this has not been demonstrated yet. Furthermore, in a seminal work Espinoza-Vergara and coworkers showed that *V. cholerae* released from EFVs exhibited colonization advantages when inoculated in the mouse digestive tract [11]. The molecular mechanisms underlying bacterial packaging into EFVs remain to be elucidated. While it is believed to be a protozoan-driven process, bacterial virulence factors appear to play an important role [10, 11, 14].

The opportunistic pathogen *Burkholderia cenocepacia* is a member of the *Burkholderia cepacia* complex (Bcc), a group of 24 closely related bacterial species known to cause life-threatening infections in cystic fibrosis (CF) and chronic granulomatous disease (CGD) patients [15, 16]. Moreover, Bcc bacteria have also been associated with nosocomial infections in immunocompromised and immunocompetent patients hospitalized for reasons unrelated to CF and CGD [17]. Currently, *B. cenocepacia* and *B. multivorans* are the Bcc species most frequently isolated from patients in Europe and North America [18]. *B. cenocepacia* is multidrug resistant and transmissible among CF patients, and causes a fatal condition known as “cepacia syndrome” which is characterized by necrotizing pneumonia and progressive respiratory failure. In particular, *B. cenocepacia* strain K56-2 is a member of the highly transmissible epidemic lineage ET12 that is responsible for most outbreaks [19]. The isolation of clonal strains of *B. cenocepacia* from both infected patients and the environment suggests that the latter is a potential reservoir for infection in the absence of nosocomial or patient-to-patient transmission [20, 21]. *B. cenocepacia* survives within mammalian macrophages [22] and human airway epithelial cells [23]. However, little is known about its interactions with non-human hosts. *In vitro, B. cenocepacia* has been shown to kill the nematode *Caenorhabditis elegans* [24], and is able to survive without replicating inside acidic vacuoles of the free-living amoeba *Acanthamoeba polyphaga* [25]. Thus, although amoebae were proposed to serve as reservoirs for Bcc bacteria in the environment [26], the role that protozoa play in the epidemiology and ecology of Bcc is currently unknown. To date, no study has investigated whether Bcc bacteria can be packaged into EFVs by protists. Hence, as ciliates and Bcc bacteria co-occur in the same ecological niches [27] and given the likely implication of protozoan EFVs in pathogen dissemination, we aimed at better understanding the interaction between ciliates and *B. cenocepacia*. By gaining insights into Burkolderia-ciliates interactions, we hope to identify potential pathways for controlling the spread of Bcc bacteria and reducing the risk of infections associated with them.

Here, we report for the first time that ciliates isolated from hospital and domestic sink drains expel EFVs laden with viable *B. cenocepacia* bacteria. Remarkably, our data demonstrates that passage through ciliates enhanced stress resistance and produced antibiotic tolerant persister cells in *B. cenocepacia*. Persisters are defined as subpopulations of bacterial cultures that transiently adopt a phenotype characterized by a non-growing state tolerant to lethal concentrations of antibiotics [28]. They are a major contributor to treatment failure and recalcitrance of chronic infections as those in CF patients. Besides stochastic generation in bacterial populations [29], environmental stress is believed to trigger persister formation as well [28, 30]. While persister cells have been identified in all major pathogens, only a few studies have investigated the switch to the persister phenotype within eukaryotic cells [31–33]. Notably, our data demonstrate that SOS activation by ROS is required for persister formation within the protozoan phagosome, further revealing that ciliate food vacuoles are genuine hot spots for the generation of antibiotic-tolerant persisters in the natural and the built environments.

## Materials and methods

### Bacterial and protozoan strains and growth conditions

The bacterial strains and plasmids used in this study are listed in Tables S1 and S2, respectively. Strains of *B. cenocepacia* and *E. coli* were grown in Luria-Bertani (LB) medium at 37 °C. When required, antibiotics were added to the following final concentrations: 100 μg/ml trimethoprim (Panreac) for *B. cenocepacia* and 50 μg/ml for *E. coli*, 150 μg/ml tetracycline (Panreac) for *B. cenocepacia* and 20 μg/ml for *E. coli*, and 40 μg/ml kanamycin (Panreac) for *E. coli*. Gentamicin (Panreac) was used at 50 μg/ml in conjugation assays to select against donor and helper *E. coli* strains.

*T. elliotti* strain 4EA was cultured at 28 °C in 2% proteose peptone (Pronadisa) supplemented with 10 μM FeCl_3_ (Sigma Aldrich) and 250 μg/ml of both streptomycin sulfate and penicillin G (Sigma Aldrich). *Colpoda* sp strain CSE36 and *Tetrahymena* sp strain T2305B2 were routinely cultured at 28 °C in PAGE’s saline buffer [34] with heat-killed inactivated *E. coli* K-12 as food source.

### Construction of reporter and mutant strains

Unmarked non-polar deletion mutants were constructed by using the approach of Flannagan and co-workers [35], which is based on the homing nuclease I-SceI. To delete the *recA* gene, the knockout plasmid pFA094 was generated by PCR amplification with Phusion HF DNA polymerase (Thermo Scientific) of the upstream and downstream flanking regions of *recA* from *B. cenocepacia* K56-2 gDNA (locus tag K562_10985, accession number NZ_CP053300) and primers FA198 to FA200, incorporating the XbaI and EcoRV restriction sites to insert them into the pGPI-SceI plasmid. Plasmid pFA094 was transferred into *B. cenocepacia* K56-2 by tri-parental mating using *E. coli* HB101 harbouring plasmid pRK2013 as helper strain. Briefly, overnight cultures of donor, helper and recipient strains were resuspended in fresh SOB medium and combined in a 3:3:1 ratio, respectively. Mating mixtures were spotted onto a nitrocellulose filter overlaid on LB agar plates and incubated at 37 °C for 16 h. Exconjugants were selected on LB medium with 100 μg/ml trimethoprim and 50 μg/ml gentamicin and screened by colony PCR. Positive clones were conjugated with *E. coli* carrying plasmid pDAI-SceI-SacB to induce a second crossover event. Tetracycline-resistant trimethoprim-sensitive exconjugants were selected and confirmed for *recA* deletion by PCR analysis. The deletion mutants were then cured from pDAI-SceI-SacB by growing in LB overnight and screening for loss of tetracycline resistance. The Δ*recA* phenotype was confirmed by its higher susceptibility to UV compared to the wild type strain K56-2 (Supplementary Material). The *B. cenocepacia* Δ*recA* mutant was complemented as follows. The coding region of *recA* was PCR amplified with primers FA219 and FA220 from gDNA of K56-2 and cloned into NdeI/XbaI sites of plasmid pSCrhaB2plus [36]. The resulting plasmid, named pFA107, was introduced into K56-2 Δ*recA* strain by tri-parental mating as described above. Transconjugants harbouring the complementing plasmid were selected on trimethoprim and gentamicin containing plates and confirmed by PCR. This strain was named as FA108.

The *lexA(Ind-)* mutant strain was obtained by gene replacement. To generate the *lexA*(S119A) mutation cassette, two fragments containing the 500 bp upstream region with the complete *lexA* CDS (locus tag K562_12117, accession number NZ_CP053300) and the 500 bp downstream region were PCR amplified with Phusion HF DNA polymerase (Thermo Scientific) from K56-2 gDNA using primers FA225-FA226 and FA227-FA228, respectively. The tetracycline cassette was PCR amplified from pDAI-SceI-SacB using primers FA223 and FA224. Then, the three fragments were PCR assembled together using the 20 bp homology regions in primers FA226 and FA227, and introduced into pGPI-SceI by restriction cloning with XbaI and EcoRI (Thermo Fisher Scientific). The point mutation *lexA*(S119A) was introduced using the Q5-Site-directed mutagenesis kit (New England Biolabs) with primers FA229 and FA230. The resulting mutant allele was verified by Sanger sequencing with primers FA236 and FA237. Plasmid carrying the *lexA*(S119A)::Tet^r^ cassette was named pFA118 and was delivered to K56-2 by triparental mating as described before. Tetracycline-resistant and trimethoprim sensitive exconjugants were selected and the presence of the mutant allele *lexA*(S119A) was confirmed by PCR and DNA sequencing. The *lexA(Ind-)* phenotype was confirmed by its higher susceptibility to UV compared to the wild type strain K56-2 (Supplementary Material).

The FA115 strain was generated by introducing pFA158 into K56-2 by triparental mating as described before and selecting for trimethoprim and gentamicin resistant exconjugants. pFA158 was generated by cloning the *dsRed2* coding sequence into NdeI/XbaI sites of plasmid pSCrhaB2plus. *dsRed2* coding sequence was PCR amplified from plasmid pUC18T-mini-Tn7T-Tp-dsRedExpress [37] with primers FA265 and FA266.

The FA161 (*PrecA-egfp)* reporter strain was generated by introducing pFA160 into K56-2 by triparental mating as described above. pFA160 was generated by replacing the S7 promoter in pMLS7eGFP [38] with the K56-2 *recA* promoter, which was previously PCR amplified with primers FA289 and FA290 from K56-2 gDNA with Phusion HF DNA polymerase (Thermo Scientific).

The oligonucleotides used in this study are listed in Table S3.

### Assessment of *B. cenocepacia* resistance to protozoan predation

Bacterial intracellular survival was assessed by quantifying *B. cenocepacia* survival in co-culture with the protozoan predator. Briefly, synchronized log-phase cultures of bacteria (OD_600_=0.3) and ciliates (3×10^5^ cells/mL) were washed and resuspended in 0.01 M Tris-HCl pH 7.5 buffer. Co-cultures were established by combining protist and bacteria (1:200 ratio) in a 96-well plate for up to 48 h at 28 °C. Every 24 h, intracellular bacteria were released by lysing protozoan cells with 1% Triton X-100 (Sigma Aldrich). The number of viable bacteria were quantified by CFU counts on LB agar plates. For each assay, at least three independent experiments were performed. Additionally, the Live/Dead BacLight Bacterial Viability kit (Themo Fisher Scientific) was used to assess the proportion of viable bacterial cells contained within EFVs. For that purpose, co-cultures, set up as described above, were incubated for 15 min with the Live/Dead BacLight Bacterial Viability solution (Thermo Scientific) according to manufacturer’s directions. Cells and EFVs were then washed twice with 0.01 M Tris-HCl pH 7.5 and observed under an Olympus CX41 fluorescence microscope using the 470–490 nm excitation/515–565 emission and 500–530 nm excitation/590–620 nm emission filter sets.

### Transmission electron microscopy

Samples from *B. cenocepacia* and *T. elliotti* co-cultures were fixed for 1 h in 0.1 M sodium cacodylate buffer pH 7.2 (TAAB Laboratories Equipment Ltd) containing 2.5% glutaraldehyde (Sigma Aldrich). Cells were then washed three times in cacodylate buffer and fixed with 0.5% OsO_4_ (TAAB Laboratories Equipment Ltd) in cacodylate buffer for 45 min at 4 °C. Fixed cells were stained for 1 h with 1% uranyl acetate (TAAB laboratories Equipment Ltd), and later dehydrated in an acetone series of increasing concentration (24%, 50%, 75% and 100%). After that, cells were embedded in Embed 812 resin (TAAB Laboratories Equipment Ltd) by following manufacturer’s directions. Ultrathin sections were obtained with microtome Ultracut (Reichert-Jung), stained with 2% uranyl acetate and lead citrate and observed in a JEM1010 transmission electron microscope at 80 kV at the Spanish National Centre for Electron Microscopy - Complutense University of Madrid.

### Detection of ROS production within protozoan phagosomes

ROS production within protozoan food vacuoles was investigated by staining cells with the ROS indicator 2’,7’-Dichlorofluorescein diacetate (DCFH-DA, Sigma Aldrich). After deacetylation by cellular esterases, DCFH turns into the green fluorescent product 2’,7’- Dichlorofluorescein (DCF) upon oxidation by intracellular ROS. Log-phase bacteria were stained with 10 µM DCFH-DA for 30 min, washed twice and resuspended in 0.01 M Tris-HCl pH 7.5. Then, stained bacteria were fed to ciliates using a 1:200 protist:bacteria ratio and transferred to a 96 well black plate (Thermo Scientific). DCF fluorescence was monitored automatically by measuring green fluorescence with 480 nm excitation and 512-552 nm emission wavelengths with a TECAN Infinite MPlex plate reader every 15 min. The fluorescence signal was normalized to OD_600_ nm. In addition, ROS production within protozoan food vacuoles was visualized by co-incubating dsRed-expressing bacteria with 10 μM DCFH-DA-stained *T. elliotti* cells. After 1 h of co-incubation at 28 °C ciliates (and ingested bacteria) were harvested, washed twice with 0.01 M Tris-HCl pH 7.5 and observed under an Olympus CX41 fluorescence microscope using the 470–490 nm excitation/515– 565 590-620 nm emission filter set.

### *PrecA-egfp* reporter assay

Co-cultures protist:bacteria were established in 0.01 M Tris-HCl pH 7.5 buffer as described above (1:200 protist:bacteria ratio) and transferred to a 96 well black plate (Thermo Scientific). GFP fluorescence was monitored automatically by measuring green fluorescence with 480 nm excitation and 512-552 nm emission wavelengths nm with a TECAN Infinite MPlex plate reader (Tecan Group Ltd) every 30 min for 20 h at 28 °C. The fluorescence signal was normalized to OD_450_.

### RNA extraction and quantitative reverse transcription PCR (qRT-PCR)

Co-cultures protist:bacteria (1:200) were established in 0.01 M Tris-HCl pH 7.5 buffer as described above and incubated at 28 °C for 24 h. At each time point, intracellular bacteria were released by lysing ciliates on ice for 20 min in 0.1% SDS (Fisher Scientific), 1% acidic phenol (Fisher Scientific), 19% ethanol in DEPC-H_2_O_2_ (Sigma Aldrich). Bacteria were then collected by centrifugation (3500 xg, 15 min) and total RNA was extracted and DNAseI-treated using the Direct-zol RNA miniprep plus kit (Zymo Research). RNA quality was verified by 1% agarose electrophoresis and quantified using Nanodrop One (Thermo Scientific). As control, total RNA was also isolated from bacteria incubated for 24 h in 0.01 M Tris-HCl pH 7.5 in the absence of ciliates by following the same procedure.

Isolated RNA was retrotranscribed with the Maxima Reverse Transcriptase (Thermo Scientific) by following manufacturer’s directions. cDNA samples were then amplified in duplicate in 96 microtiter plates (4titude^®^). Real time PCR was carried out in 20 µl reaction mixtures containing 1x TB Green Premix Ex Taq (Takara), 1x ROX reference dye (Takara), 300 nM of each forward and reverse primer (Integrated DNA Technologies) and 2 µl (25 ng) of the template cDNA. Samples were amplified in a iQ5 real-time PCR (Bio-Rad) using the following thermal cycling protocol: 5 min at 95 °C, followed by 40 cycles of 30 sec at 95 °C, 30 sec at 55 °C and 20 sec at 72 °C; and a final step of 1 min at 95 °C and 1 min at 55 °C. The specificity of each primer pair was confirmed by melting curve analysis. The fold change in mRNA expression levels was calculated with the 2^-ΔΔCt^ method [39] and normalized against the expression level of the housekeeping gene *gyrA* (locus tag K562_12858, accession number NZ_CP053300). Non-template controls (NTC) and RT minus controls were negative.

### Assessment of stress resistance of *B. cenocepacia* recovered from protists and persister assays

Co-cultures protist:bacteria (ratio 1:200) were established in 0.01 M Tris-HCl pH 7.5 buffer as described above. As control, the same number of log-phase bacteria was incubated buffer in the absence of ciliates. After 24 h incubation, EFVs were collected from co-cultures and washed twice with 0.01 M Tris-HCl pH 7.5 buffer. Bacteria were then released by adding 1% Triton-X100 and pelleted by centrifugation at 3500 xg for 5 min. Released bacteria were washed twice with 0.01 M Tris-HCl pH 7.5 buffer. In parallel, control bacteria incubated in 0.01 M Tris-HCl pH 7.5 buffer for 24 h were harvested, exposed to 1% Triton-X100 and washed twice with 0.01 M Tris-HCl pH 7.5 buffer. At this point both bacteria released from EFVs and control bacteria were subjected to different stress conditions (0.02% H_2_O_2_, starvation and desiccation) for the indicated time intervals. At selected time points bacterial suspensions were subsequently serially diluted 1:10 in 0.9% NaCl and plated on LB agar plates. The number of surviving bacteria was estimated by CFU counts. For each assay three independent experiments were performed. In desiccation resistance assays, bacteria were spotted onto cellulose filter disks and dried at 30 °C for up to 24 h, then rehydrated in 0.9% NaCl for 30 min and spread on LB agar plates to allow for CFU counts.

For persister assays, bacteria (released from EFVs and controls prepared as described above) at 10^7^ CFUs/mL were incubated in LB buffered medium supplemented or not (control) with 40-80 μg/mL (10-20x MIC) ciprofloxacin (Sigma Aldrich) or 320 μg/mL (10x MIC) meropenem (Sigma Aldrich) for 20 h at 37 °C. At given time points, bacteria were collected, resuspended and washed in 10 mM MgSO_4_ (Sigma Aldrich) to inactivate the residual antibiotic. The bacterial suspensions were then serially diluted in 0.9% NaCl and plated on LB agar plates to quantify CFUs. Surviving bacteria were harvested from the plates, grown overnight in fresh LB and used to perform a follow-up persister assay.

### Paraquat protection assay

Bacteria from synchronized log-phase cultures (OD_600_ = 0.3) were incubated in LB buffered medium supplemented or not (control) with 1 mM Paraquat (Sigma Aldrich) for 30 min at 37 °C in a 12-well polystyrene plate. The number of viable bacteria after treatment (time=0) was estimated by plating serial dilutions on LB agar plates and CFU counts. Then, 40 μg/mL ciprofloxacin was added to each well and bacteria were incubated with the antibiotic for up to 20 h at 37 °C. At given time points, bacteria were collected from each well, washed in 10 mM MgSO_4_ to inactivate the residual antibiotic, and plated on LB agar to estimate the number of survivors by CFU counting. For each time point, the percentage of survival after ciprofloxacin challenge was determined by comparing number of survivors after antibiotic exposure to survivors of the untreated group at the corresponding time point.

### NOX inhibition assay

For NOX inhibition assays, *T. elliotti* cells (5×10^5^ cell/mL) were pre-treated with 10 μM VAS2870 (Cayman Chemicals) for 1 h at 28 °C before adding the bacteria. Then, bacteria from synchronized log-phase cultures (OD_600_ = 0.3) were added to pre-treated and untreated (control) ciliates and incubated for 3 h at 28 °C. Afterwards, ciliates were lysed with 1% Triton X-100 and the released bacteria were washed and incubated at 37 °C in fresh LB buffered medium supplemented with 40 μg/mL (10x MIC) ciprofloxacin in a 24-well plate. Three hours later, bacteria were collected from each well, washed in 10 mM MgSO_4_ and plated on LB agar to estimate the number of survivors by CFU counting. The percentage of survival after ciprofloxacin challenge was determined by comparing survivors after 3 h of antibiotic exposure to survivors of the corresponding untreated group at the same 3 h interval.

The *PrecA:egfp* reporter assay was performed by incubating the *B. cenocepacia* strain FA161 with ciliates pre-treated for 1 h with 10 μM VAS2870 and untreated ciliates (control) in a 24-well plate. After 3.5 h of co-incubation with bacteria, ciliates were harvested and lysed with 1% Triton-X100 to release ingested bacteria from food vacuoles. Aliquots (150 μl) from each sample were transferred to a 96 well black plate (Thermo Scientific) and GFP fluorescence was measured with a TECAN Infinite MPlex plate reader (480 nm excitation/522-552 nm emission filter). Samples were measured in triplicate. VAS2870- treated ciliates incubated in the absence of bacteria, and bacteria incubated without ciliates were used as fluorescence controls. Three independent experiments were performed.

### Statistical analysis

Statistical analyses were performed with GraphPad Prism v.8 (GraphPad Software). Statistical differences were determined using unpaired Student’s t-test when comparing two conditions and one-way analysis of variance (ANOVA) with Dunnett’s post-test when comparing three or more conditions. Details of statistical analysis performed for each set of data are shown in the figure captions.

## Results

### *B. cenocepacia* bacteria resist intracellular digestion in ciliates and are packaged into EFVs with enhanced tolerance towards stress and antibiotics

Survival assays were performed to determine whether *B. cenocepacia* K56-2 was able to evade intracellular digestion and/or multiply within different ciliates. As shown in Figure 1 A, *B. cenocepacia* survived predation although we observed different degree of grazing resistance depending on the ciliate predator. *Colpoda* sp CSE36 (wild strain isolated from domestic sink drain) caused a greater reduction (2 log_10_) in the K56-2 population, whereas *T. ellioti* 4EA (laboratory strain) and *Tetrahymena* sp T2305B2 (wild strain isolated from hospital sink drain) reduced it in less than 1 log_10_ (Figure 1, A). Accordingly, epifluorescence microscopy after Syto9/PI staining (Figure 1, B) and TEM (Figure 1, C-G) confirmed that most *B. cenocepacia* cells avoided intracellular digestion and viable bacteria, sometimes dividing, were packaged in membrane-bound vesicles that were eventually expelled by ciliates into the extracellular medium within 3 h post-feeding, similar to previously reported *L. pneumophila* or *V. cholerae* laden EFVs [9, 11]. Similar results were found when we observed EFVs produced by a wild *Tetrahymena* sp T2305B2 isolated from hospital sink drain (Figure S1).

**Figure 1.**
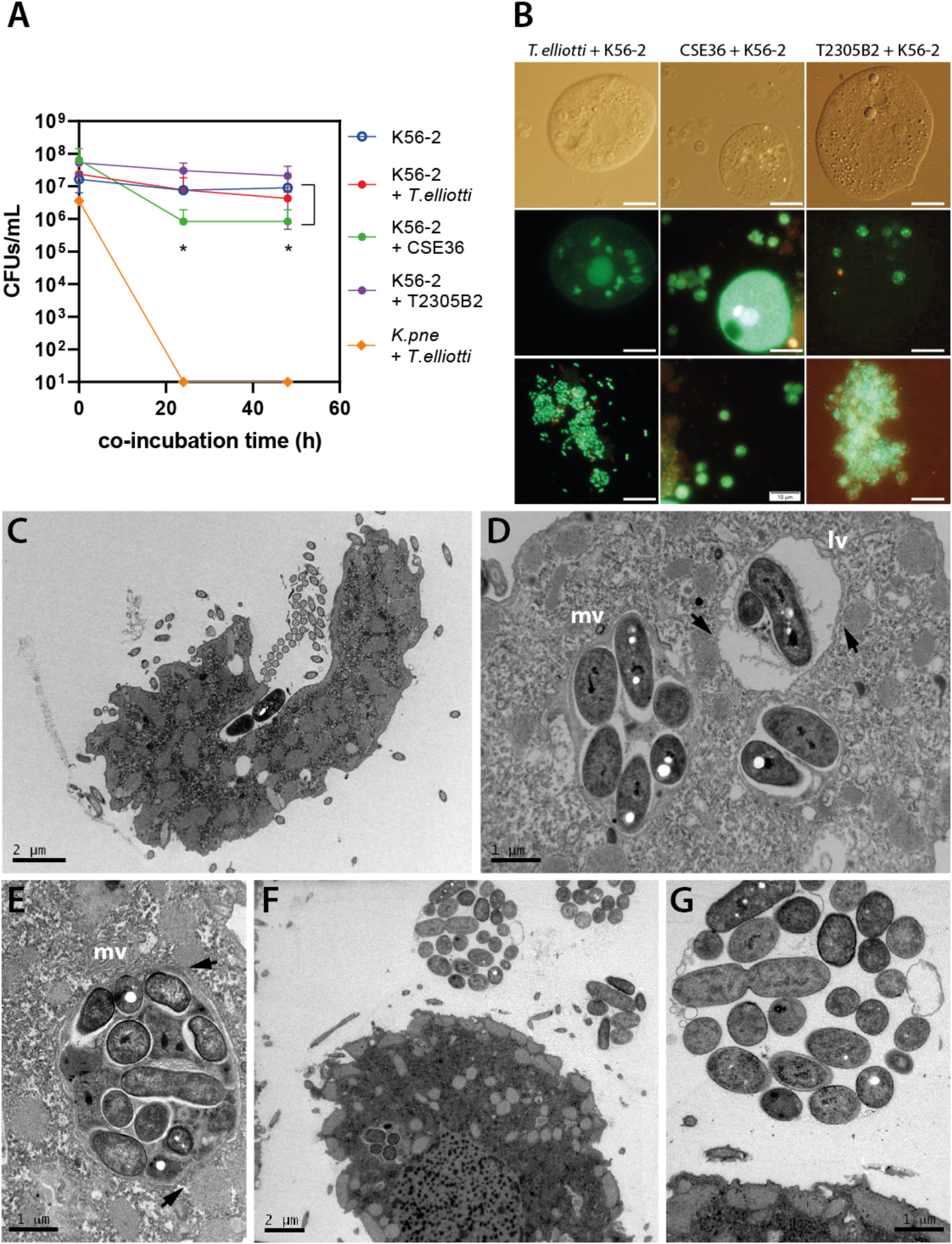
Grazing resistance of *B. cenocepacia* K56-2 and packaging of viable bacteria within EFVs by ciliates. **(A)** *B. cenocepacia* K56-2 survival under predation by wild (CSE36 and T2305B2) and domesticated (*T. elliotti*) ciliates. The number of surviving bacteria incubated in the absence (empty circles) or presence (filled circles) of ciliates was quantified by CFU counts after 0-48 h of co-incubation. The star (*) indicates significant differences (p < 0.05) between K56-2+CSE36 vs K56-2 samples according the unpaired Student’s t-test. **(B)** Viability of ingested *B. cenocepacia* K56-2 and bacteria packaged inside EFVs produced by *T. elliotti* 4EA (pictures on the left), *Colpoda* sp CS36 (center) and *Tetrahymena* sp T2305B2 (right). Aliquots of the bacteria:ciliate co-cultures were stained after 24 h of co-incubation with the Live/Dead BacLight solution and observed with an epifluorescence microscope to visualize intact (green, Syto9-stained) or membrane-damaged (red, PI-stained) bacterial cells within intracellular food vacuoles and EFVs. White scale bar represents 10 μm. **(C-G)** TEM micrographs of *T. elliotti* grazing on *B. cenocepacia* K56-2. Bacteria being engulfed within a nascent food vacuole at the ciliated cytopharynx (C). Most K56-2 bacteria survived within mid-stage and late-stage food vacuoles (D, E) and were eventually packaged into EFVs and expelled by ciliates (F, G). Mid-stage (mv) food vacuoles are characterized by a membrane that closely molds to the shape of enclosed bacteria. In contrast, late-stage (lv) vacuoles are spherical and exhibit a distinctive space or halo between the enclosed bacteria and the smooth vacuole’s membrane. Arrowheads point at mitochondria located near the food vacuoles. Dividing bacteria were observed within EFVs (G).

Since passage through ciliates has been shown to increase environmental fitness in several grazing resistant bacteria [11–13], we assessed the stress resistance of K56-2 cells recovered from *T. elliotti* EFVs. Because we could not establish axenic cultures of undomesticated ciliates isolated from sink drain biofilms, we performed these experiments using *T. elliotti* 4EA as predator to avoid possible interferences with the indigenous bacterial species carried by the wild protists. We believe *T. elliotti* 4EA is a good surrogate model since we have isolated *Tetrahymena* species such as strain T2305B2 from hospital and domestic sink drains such (Amaro, unpublished data). Thus, packaged K56-2 bacteria from overnight produced *T. elliotti* EFVs were released with 1% Triton X-100 and subsequently exposed to different stress conditions: H_2_O_2_, starvation, desiccation, and antibiotics (ciprofloxacin, meropenem, tobramycin). As control, K56-2 cells incubated overnight in 0.01 M Tris-HCl buffer pH 7.5 in the absence of ciliates were exposed to same conditions. As shown in Figure 2A, bacteria recovered from EFVs exhibited a higher resistance towards H_2_O_2_ exposure, surviving (0.5 log_10_ reduction, 26 % survival after 60 min) at H_2_O_2_ doses that eradicated control bacteria in 60 min (6 log_10_ reduction, < 0.0001 % survival). Additionally, EFV-released K56-2 bacteria survived desiccation better (3 orders of magnitude higher after 6 h, 1 order of magnitude higher after 24 h) than control K56-2 did (Figure 2 B). However, we did not observe any protective effect towards starvation, as similar survival rates were observed for control and EFV-released bacteria (data not shown). Remarkably, bacteria recovered from EFVs exhibited a higher tolerance towards high concentrations (10-20x MIC) of different classes of antibiotics. As shown in Figure 2 C-D (and Figure 5 E), passage through *T. elliotti* produced a 10 and >100-fold increase in the proportion of bacteria tolerant to meropenem (a carbapenem, beta-lactam that blocks peptidoglycan biosynthesis) and ciprofloxacin (a quinolone, DNA-gyrase inhibitor), respectively. Time-kill curves performed in the presence of high concentrations (10x MIC) of these antibiotics alone revealed a biphasic killing curve, suggesting that antibiotic-tolerant bacteria recovered from EFVs were persisters (Figure 2 D). Accordingly, the progeny of bacteria that survived antibiotic challenge were as susceptible as the initial inoculum and simil biphasic killing curves were obtained (data not shown), indicating that they were *bona fide* persisters. However, bacteria released from protozoan EFVs were killed by >4x MIC tobramycin (data not shown), indicating that persisters generated within *T. elliotti* food vacuoles were susceptible to this aminoglycoside. Consistent with our results, susceptibility to aminoglycosides in persisters that are tolerant to quinolones and beta-lactams has been thoroughly documented [40–44]. Overall, these results show that successful passage through ciliates confer enhanced resistance to oxidative stress and desiccation as well as some of the antibiotics that are prescribed to treat *B. cenocepacia* infections [45, 46], which could favor pathogen dissemination in the environment (natural or man-made) and colonization of the human host.

**Figure 2.**
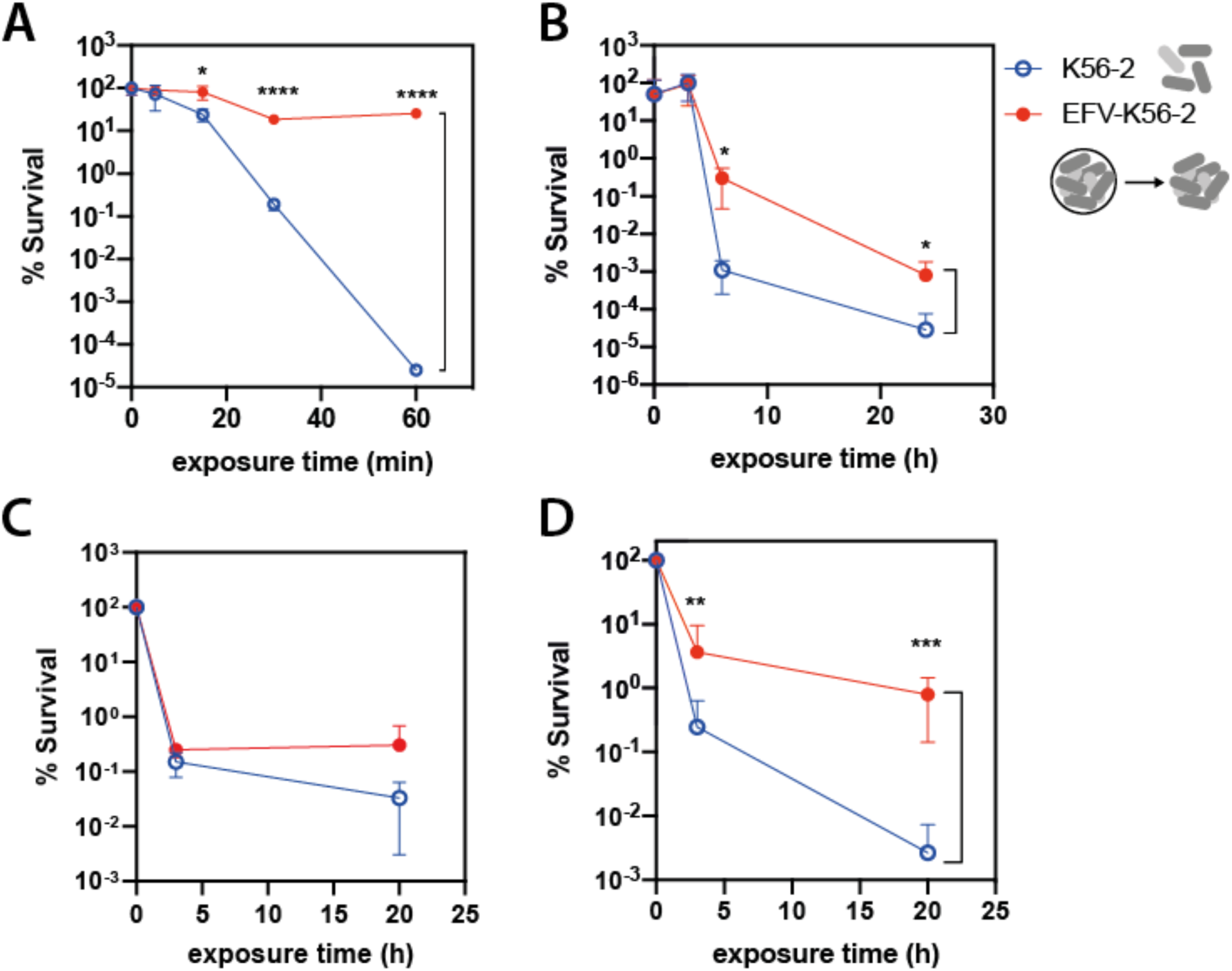
Increased stress resistance and antibiotic tolerance of *B. cenocepacia* K56-2 after passing through *T. elliotti*. **(A-D)** Survival rate of K56-2 cells released from EFVs (filled circles) and control planktonic K56-2 cells (empty circles) subjected to 0.02% H_2_O_2_ (A), desiccation (B), 320 μg/mL (10x MIC) meropenem (C) and 40 μg/mL (10x MIC) ciprofloxacin (D). At selected time points, bacteria were plated on LB and surviving bacteria were quantified by CFU counts. Data shown correspond to the average value from three independent biological replicates. Significant differences (*p < 0.05, **p < 0.01, ***p < 0.001, ****p < 0.0001) between EFV-K56-2 and K56-2 (control) for each time point were determined using unpaired Student’s t-test.

### ROS generated within *T. elliotti* phagosomes trigger the SOS response in *B. cenocepacia* and promote antibiotic tolerance

Since we had observed a greater protection against ciprofloxacin than meropenem, we focused on investigating the underlying mechanism conferring protection towards ciprofloxacin during passage through ciliates. Because bacterivorous protists and macrophages challenge ingested bacteria with an oxidative burst in the phagosome [47], we first examined ROS production within the food vacuoles of *T. elliotti* grazing on K56-2 cells pre-stained with DCFH-DA, a well-known ROS probe which is oxidized by ROS to the green fluorescent 2′,7′-dichlorofluorescein (DCF) [48]. As shown in Figure 3 A, DCFHDA-stained K56-2 cells exhibited a dramatic increase in green fluorescence 15 min upon feeding them to *T. elliotti*, similar to that shown by stained bacteria exposed 0.02 % H_2_O_2_, thus suggesting ROS production and subsequent oxidation of DCFH-DA to fluorescent DCF. Additionally, ROS generation within *T. elliotti* phagosomes was corroborated in a different assay by feeding dsRed-labeled *B. cenocepacia* K56-2 to DCFHDA-stained *T. elliotti*. As shown in Figure 3 B, bright green fluorescence produced by oxidized DCF co-localized with dsRed-expressing bacteria inside *T. elliotti* food vacuoles, confirming the exposure of K56-2 to ROS within protozoan phagosomes.

**Figure 3.**
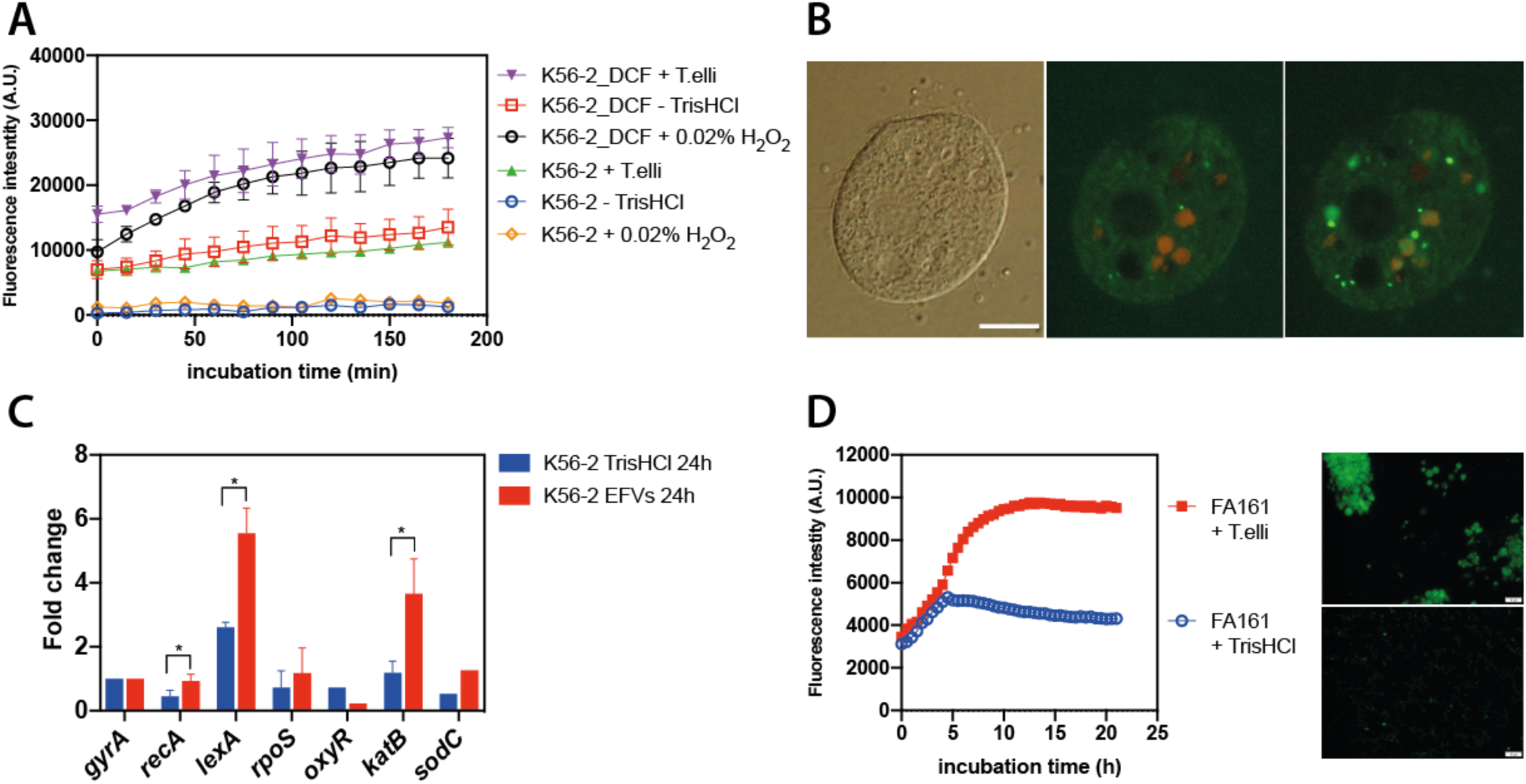
ROS production and upregulation of *B. cenocepacia* stress-related genes in protozoan phagosomes. **(A)** Quantification of ROS production as green fluorescence emitted by DCFHDA-stained K56-2 bacteria ingested by *T. elliotti*. Bacteria were stained with DCFH-DA for 30 min, washed and fed to *T. elliotti*. Green fluorescence produced by oxidized DCF was monitored for 180 min. A dramatic increase in the production of green fluorescence was detected for DCF-stained K56-2 cells that were incubated in the presence of *T. elliotti* (purple line) or 0.02% H_2_O_2_ (orange line, positive control). **(B)** Co-localization of ROS and bacteria in protozoan food vacuoles was demonstrated by feeding dsRed-expressing K56-2 bacteria to DCFHDA-stained *T. elliotti* (after 3h co-incubation). **(C)** Upregulation of SOS response and oxidative stress related genes in *B. cenocepacia* K56-2 recovered from protozoan food vacuoles (at 24h post-feeding). Significant differences (*p < 0.05) were determined with the unpaired Student’s t-test. **(D)** Time-course quantification of GFP fluorescence emitted by the reporter strain *B. cenocepacia* FA161 (PrecA:eGFP) incubated with (red line, top picture) or without (blue line, bottom picture) *T. elliotti*. The dramatic increase in GFP fluorescence detected in the presence of ciliates suggests the upregulation of the SOS response in ingested bacteria. Scale bar represents 10 μm.

Furthermore, considering ROS production within protozoan food vacuoles, and the known association between ciprofloxacin tolerance and the SOS response in *E. coli* [49], we hypothesized that oxidative radicals encountered in the protozoan phagosome may promote persister switch by activating the SOS response in *B. cenocepacia*. To investigate that, we first confirmed SOS upregulation within protozoan food vacuoles. We used qRT-PCR to quantify the expression levels of SOS and oxidative stress related genes in *B. cenocepacia* cells collected from *T. elliotti* EFVs. Compared to bacteria incubated in the absence of ciliates, the relative expression of *recA*, *lexA*, *katB* and *sodC* was >2-fold higher in bacteria isolated from EFVs (Figure 3 C). Therefore, upregulation of genes encoding catalase KatB and superoxide dismutase SodC, two enzymes involved in detoxification of H_2_O_2_ and O_2.-_, respectively, confirmed that bacteria were exposed to ROS within *T. elliotti* food vacuoles. Likewise, overexpression of genes encoding the two SOS regulators *recA* and *lexA* suggested activation of the SOS response. Upon DNA damage, activated RecA protein triggers the SOS response by inducing self-cleavage of the autoregulated repressor LexA, thus releasing the transcription of LexA-regulated genes (*recA*, *lexA* and *uvrA*, among others), increasing their mRNA levels in the cell [50]. Supporting our data, similar results were obtained for *B. cepacia* cells recovered from *T. elliotti* EFVs (Figure S2). Additionally, upregulation of *recA* in ingested bacteria was confirmed by feeding the reporter strain *B. cenocepacia* FA161 (which expresses *gfp* under the control of the *recA* promoter) to *T. elliotti*. A dramatic increase in GFP fluorescence was observed upon 4 h of co-incubation compared to the strain incubated in the absence of ciliates (Figure 3 D). In addition, GFP fluorescent bacteria were observed inside intracellular food vacuoles and EFVs as well. Indeed, bacteria exhibited a brighter GFP fluorescence upon passage through *T. elliotti* when compared to bacteria incubated in the absence of ciliates (Figure 3 D). Therefore, these results indicated that conditions encountered within the protozoan food vacuoles trigger the SOS response in *B. cenocepacia*.

We then tested whether ROS exposure promotes antibiotic tolerance in *B. cenocepacia*. To assess that, we pre-treated log-phase bacterial cultures with sub-lethal doses of the ROS-generating agent paraquat (PQ) for 30 min prior to exposure to high concentrations (10x MIC) of ciprofloxacin. PQ is a redox-cycling compound that produces superoxide and subsequently H_2_O_2_, generating an oxidative stress environment resembling that encountered during respiratory burst in the phagosome [33]. PQ pretreatment did not cause bacterial death (Figure S3), but increased the percentage of bacteria (persisters) that survived 3 and 20 h exposure to ciprofloxacin by 3 and 2 orders of magnitude (log_10_), respectively (Figure 4 A). Again, we observed the characteristic biphasic killing curve, suggesting that the surviving bacteria were indeed persisters. Then, given that a ROS challenge induced persister formation in LB medium, we investigated if blocking ROS production in the protozoan phagosome would diminish the percentage of persisters recovered from *T. elliotti*. To assess that, we pretreated ciliates with the NOX inhibitor 10 μM VAS2870 for 1 h prior feeding them with K56-2 bacteria. After 3 h, we lysed the ciliates and exposed the released bacteria to 40 μg/mL (10x MIC) ciprofloxacin for 3 h. Importantly, addition of the NOX inhibitor VAS2870 to ciliates significantly decreased by 10-fold the percentage of intracellular persisters recovered (Figure 4 B), supporting that an oxidative burst promotes persister switch within protozoan food vacuoles. In line with this, we confirmed that protozoan NOX contributes to *B. cenocepacia* SOS activation by feeding the SOS-reporter strain FA161 (*PrecA:egfp*) to ciliates pretreated with VAS2870 for 1 h. Untreated ciliates served as control. After 3.5 h of co-incubation with bacteria, ciliates were lysed with 1% Triton-X100 and GFP fluorescence of released bacteria was measured. As shown in Figure 4 C bacteria collected from non-treated ciliates exhibited 5-fold higher GFP fluorescence than those recovered from VAS2870-treated ciliates, indicating that NOX inhibition results in lower activation of the SOS response.

**Figure 4.**
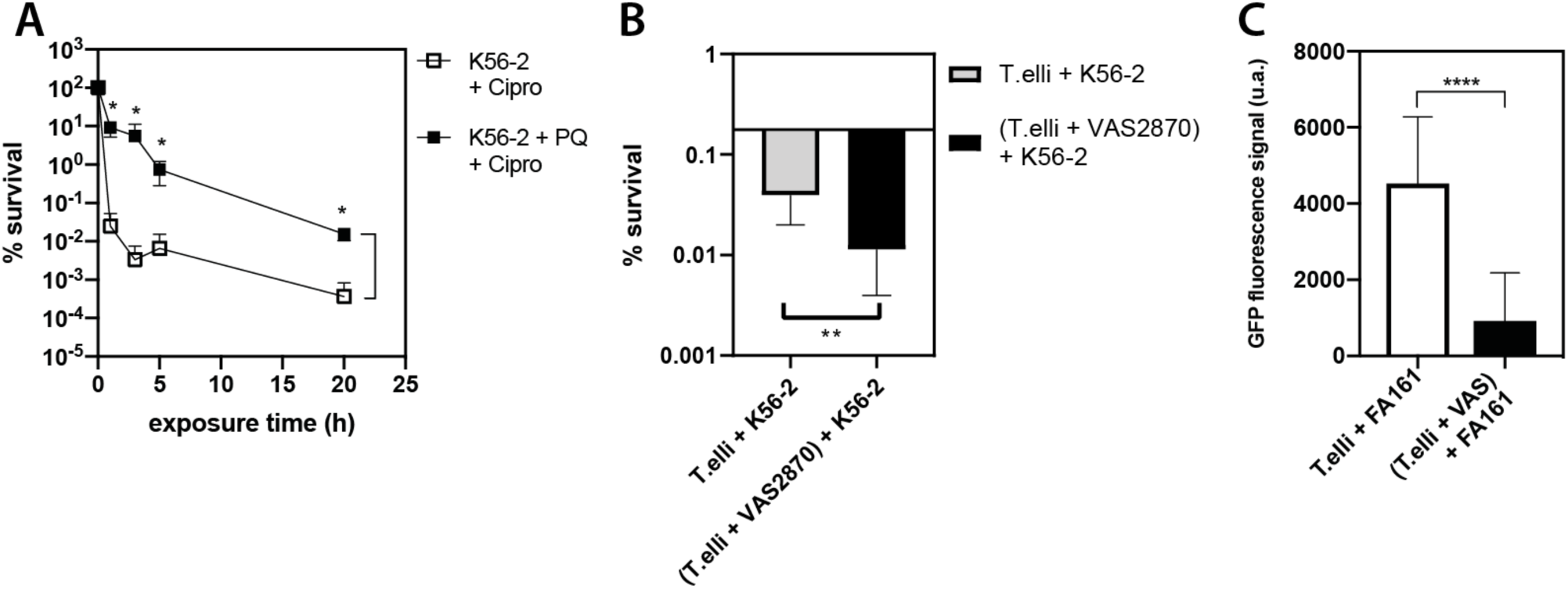
Protist-produced ROS induce persister switch in *B. cenocepacia*. **(A)** Time killing assay for log-phase K56-2 bacteria exposed to 40 μg/mL (10x MIC) ciprofloxacin in LB after being pre-treated with paraquat (PQ, filled black squares) or not (control, empty squares) for 30 min before adding the antibiotic. Sublethal pre-challenge with PQ in LB enhanced antibiotic tolerance in planktonic *B. cenocepacia* cells. The antibiotic caused a bi-phasic killing curve indicating the presence of persisters tolerant to ciprofloxacin. **(B)** Formation of *B. cenocepacia* persisters within *T. elliotti* food vacuoles. Survival percentage of K56-2 collected from digestive vacuoles from ciliates that were pretreated for 1 h with 10 μM VAS2870 (black bar) or not (grey bar) prior feeding with bacteria. Bacteria were added to untreated (control) or VAS2870-treated ciliates and incubated for 3 h. Then, intracellular bacteria were released from food vacuoles with 1% Triton-X100 and exposed to ciprofloxacin (10x MIC) in LB for 3 h at 37 °C. The percentage of bacteria that survived antibiotic (persisters) was estimated by CFU counts after plating on LB agar. Inhibiting *T. elliotti* ROS production with the NOX inhibitor VAS2870 caused a 10-fold decrease in the percentage of persisters tolerant to ciprofloxacin, compared to persisters recovered from untreated ciliates. **(C)** GFP fluorescence of internalized SOS-reporter strain FA161 (*PrecA:egfp*) recovered from untreated ciliates (control, white bar) or those pretreated with VAS2870 for 1 h (black bar) prior feeding. After 3.5 h of co-incubation with bacteria, ciliates were harvested, lysed with 1% Triton-X100 and GFP fluorescence of released bacteria was measured. Bacteria collected from untreated ciliates exhibited 5-fold higher GFP fluorescence than those recovered from VAS2870-treated ciliates, confirming than NOX contributes to SOS activation within protozoan food vacuoles. Represented data correspond to the average of three independent experiments. Statistical significance (*p < 0.05, **p < 0.01, ****p < 0.001) was determined by using the unpaired t student’s test.

### The SOS response is involved in *B. cenocepacia* intracellular survival and enhanced antibiotic/stress tolerance of expelled bacteria

To confirm the role of the SOS response in persister formation within protozoan food vacuoles we constructed the two SOS-deficient mutant strains Δ*recA* and *lexA(Ind-)*. We fed them to ciliates and monitored their intracellular survival and enhanced stress resistance/antibiotic tolerance after passage through ciliates. RecA (inducer) and LexA (repressor) are the master regulators of the SOS response [51]. Strain Δ*recA* expresses a non-functional truncated RecA protein, whereas strain *lexA(Ind-)* carries a LexA variant resistant to RecA cleavage, thus preventing induction of SOS response while maintaining RecA activity. The SOS-deficient phenotype of each strain was confirmed by assessing their higher susceptibility to UV exposure compared to the parental wild type strain K56-2 (Figure S4, B). Additionally, qRT-PCR assays corroborated the lack of SOS upregulation when both mutant strains were exposed to mitomycin C, a well-known inducer of the SOS response (Figure S4, C). Notably, we observed that *B. cenocepacia* intracellular survival was significantly impaired by mutations blocking the bacterial SOS response (Figure 5, A-B), suggesting that bacteria indeed face DNA damage within protozoan food vacuoles. Only 1.6% and 3.1% of the ι1*recA* and *lexA(Ind-)* populations, respectively, survived grazing by *T. elliotti*, whereas the wild-type K56-2 exhibited a 73% survival rate (Figure 5, B). The higher survival rate of strain *lexA(Ind-)* compared to ι1*recA* suggests that RecA-mediated homologous recombination likely contributes to DNA repair in *B. cenocepacia* during passage through ciliates. Live/Dead BacLight staining corroborated impaired survival for SOS-deficient mutants, as the EFVs produced by ciliates grazing on either ι1*recA* or *lexA(Ind-)* were filled with many membrane-damaged bacteria (PI-positive cells), whereas EFVs produced by ciliates grazing on the WT K56-2 strain contained mostly (>95 %) membrane intact cells (Syto9-positive and PI-negative cells, Figure 5 A). Furthermore, EFV-released SOS-mutants that survived intracellular digestion showed similar resistance to H_2_O_2_ and desiccation as the parental strain K56-2 incubated in Tris-HCl buffer (Figure 5, C-D). Likewise, the percentage of persisters tolerant to ciprofloxacin did not increase during passage through ciliates when the *lexA(Ind-)* strain was used (Figure 5, E), as the percentage of *lexA(Ind-)* persisters wihin EFVs was similar to that of the *lexA(Ind-)* strain incubated in buffer. Because the ι1*recA* mutant exhibited a stronger defect in intracellular survival compared to *lexA(Ind-),* we only evaluated persister formation during passage through ciliates with the *lexA(Ind-)* strain. Overall, these results indicate that SOS activation contributes to stress adaptation and persister formation within protozoan food vacuoles. Supporting these results, a Δ*surA* mutant strain with has higher susceptibility to ciprofloxacin (MIC=2 μg/mL) than K56-2 (MIC=4 μg/mL) and similar to *lexA(Ind-)* (MIC=2 μg/mL) but it retains an intact SOS response (Figure S5), increased its tolerance to ciprofloxacin after passage through ciliates (Figure 5, F). Therefore, the absence of a functional SOS response is likely the reason that *lexA(Ind-)* mutant did not increase its tolerance to this antibiotic after passage through ciliates.

**Figure 5.**
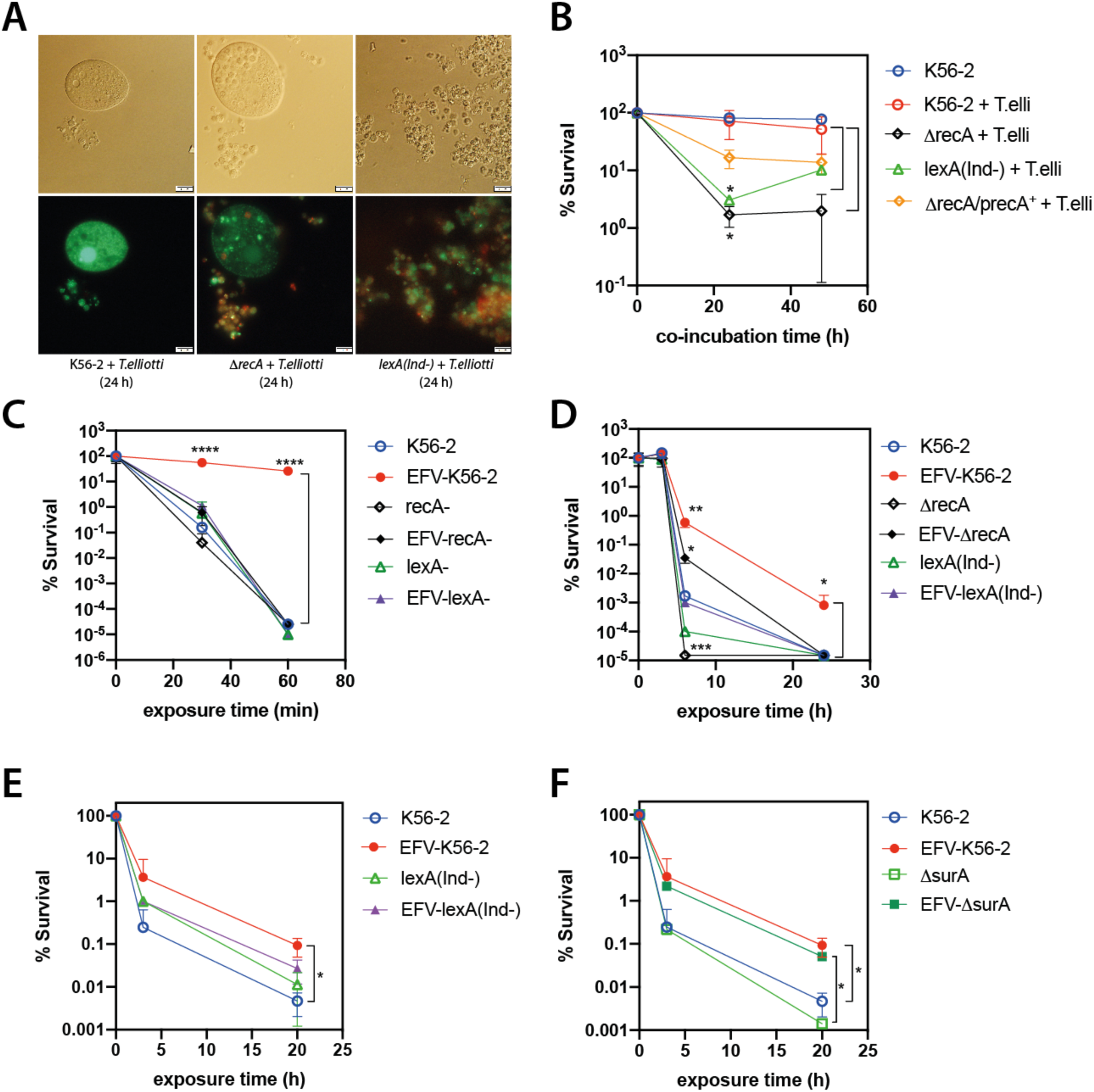
Impaired intracellular survival and stress resistance/antibiotic tolerance of WT and SOS-deficient mutant strains of *B. cenocepacia* K56-2 after passage through ciliates. **(A)** Fluorescence microscopy images of EFV-packaged bacteria stained with Syto9 and PI after 24 h of co-incubation with *T. elliotti*. EFVs produced by *T. elliotti* grazing on the SOS-deficient mutants Δ*recA* or *lexA(Ind-)* contained a higher proportion of membrane-damage bacteria (PI-positive, red fluorescence) than EFVs produced by ciliates grazing on wild-type K56-2 (EFVs filled with green bacteria, Syto9-poisitive, PI-negative). This result indicates that Δ*recA* or *lexA(Ind-)* mutations compromise *B. cenocepacia* ability to survive intracellular digestion within ciliates. **(B)** Bacterial survival under predation by *T. elliotti*. Ciliates were incubated with bacteria (ratio 1:200 protist:bacteria) in buffer. At indicated time points, ciliates were lysed with 1% Triton-X100 and surviving bacteria were enumerated by CFU counts on LB agar plates. Again, mutations in the SOS response regulators caused impaired intracellular survival within ciliates compared to the wild-type strain K56-2. Complementation of the *recA* gene in Δ*recA* strain restored intracellular survival to wild-type levels. **(C-F)** Survival rate of bacteria released from EFVs (filled symbols) and control planktonic bacteria (empty symbols) subjected to 0.02% H_2_O_2_ (C), desiccation (D), or 80 μg/mL (20x MIC) ciprofloxacin (E-F). At indicated time points, bacteria were plated on LB agar and survivors were quantified by CFU counts. Mutations in the SOS regulators decreased resistance to H_2_O_2_ and desiccation, as well as the percentage of persisters tolerant to ciprofloxacin formed within protozoan food vacuoles compared to wild-type strain (E). In contrast, the mutant Δ*surA* exhibited a tolerance to ciprofloxacin similar to that of the wild-type strain K56-2 after passage through ciliates (similar percentage of persisters) (F). Data represent the average of at least three independent biological replicates. Statistical significance (*p < 0.05, **p < 0.01, ****p < 0.001) was determined with the unpaired t student’s test.

## Discussion

It has been proposed that the induction of stress responses within protozoan food vacuoles helps bacteria to adapt themselves to the same stress conditions that they will encounter in the human host [52]. Our study evidences that *B. cenocepacia* survives phagocytosis by ciliates found in the natural and built environments, and viable bacteria are expelled packaged in fecal pellets (or EFVs) with enhanced tolerance towards harsh conditions such as desiccation, oxidative stress and antibiotics, which may significantly contribute to the persistence and transmission of this opportunistic pathogen. We show that genes encoding antioxidant enzymes are upregulated during passage through ciliates increasing bacterial resistance to ROS that could help bacteria to survive desiccation, oxidizing sanitizers or host defenses. Whereas previous studies demonstrated that packaging of some bacterial pathogens (such as *L. pneumophila, V. cholerae*, *Salmonella enterica*) into protozoan fecal pellets protects them from stress and antibiotics [11–13], we have identified that the oxidative burst generated in the protozoan phagosome promotes stress resistance and persister switch by triggering the SOS response in *B. cenocepacia*.

In the last decade, different mechanisms contributing to persister formation have been identified. These include activation of toxin-antitoxin (TA) systems, the stringent response and (p)ppGpp signaling, RpoS-mediated general stress response and the SOS response [44, 53–55]. While *rpoS* exhibited similar expression levels in K56-2 collected from EFVs or incubated in Tris-HCl buffer, *recA* and *lexA* were significantly upregulated, suggesting activation of the SOS response during passage through ciliates. Accordingly, the *PrecA-gfp* transcriptional fusion was strongly induced in *B. cenocepacia* encased within *T. elliotti* food vacuoles. In line with this, an RNA-seq study carried out in our lab did not detect upregulation of *rpoS, relA* and genes encoding known TA systems in K56-2 bacteria collected from *T. elliotti* EFVs [56]. Therefore, our data suggest that RpoS-stress response, (p)ppGpp signaling and TA systems may not be involved in formation of persisters tolerant to ciprofloxacin within ciliate food vacuoles, while the SOS response appears to play a crucial role.

Previous studies have shown that the phagosomal environment of amoebae and macrophages favors persister switch in *L. pneumophila* in a process controlled by the Legionella quorum system (Lqs) [31]. Likewise, macrophages have been found to induce antibiotic tolerance of internalized *Salmonella* serovar Typhimurium, *Mycobacterium tuberculosis* and *Staphylococcus aureus* [33, 57, 58], suggesting that persister switch may be a widespread phenomenon in intracellular bacteria surviving digestion within the phagosome. ROS are crucial components of the bactericidal repertoire of macrophages and bacterivorous protists. During oxidative burst, the membrane bound NADPH oxidase (NOX) complex generates ROS (O_2.-_ and H_2_O_2_) in the phagosomal lumen. Genes encoding NOX homologs have been identified in the genome of *T. elliotti* and other *Tetrahymena* species [59] and our data demonstrate that *B. cenocepacia* is exposed to ROS within *T. elliotti* food vacuoles. Because inhibition of the protozoan NOX complex decreased by 10-fold the number of intracellular persisters tolerant to ciprofloxacin, ROS produced in the protozoan phagosome are likely involved in persister formation within *T. elliotti* food vacuoles. In line with this, it has been recently shown that phagosomal ROS generated by macrophages induce antibiotic tolerance in intracellular *S. aureus* [33]. ROS may inhibit bacterial growth, which has been associated with antibiotic tolerance [60]. However, recent studies have shown that the induction of antibiotic tolerance cannot be solely attributed to growth inhibition by ROS [33]. Besides, we believe that growth arrest alone was not sufficient to induce ciprofloxacin tolerance in ingested K56-2 bacteria since starving bacteria incubated in Tris-HCl buffer for 24 h were rapidly killed by this antibiotic. Although persisters were initially defined as dormant cells in which essential targets that antibiotic corrupt are not active, recent studies have called this into question [28, 41], particularly within intracellular niches where bacteria must maintain their metabolic activity to withstand stressors imposed by the eukaryotic host cell [61, 62]. Moreover, persistence cannot be solely explained by growth arrest and reduced target activity for quinolones because they target topoisomerases and non-growing cells still possess transcriptional activity [44, 63]. Therefore, since both inhibition of *T. elliotti* NOX complex and mutations that abolished the SOS response significantly decreased the percentage of *B. cenoceapcia* persisters within EFVs, we propose that the SOS response triggered by oxidative burst leads to persister formation within *T. elliotti* food vacuoles. The fact that we observed lower upregulation of P*recA:egfp* in ingested bacteria when ciliates were pretreated with the NOX inhibitor supports SOS activation by phagosomal ROS. Besides, this finding is consistent with studies from other laboratories showing that activation of the SOS response confers tolerance to fluoroquinolones and beta-lactams (ampicillin) in *E. coli* and *S. aureus* [49, 64, 65]. Beyond a DNA repair process, the SOS response is now considered as a powerful strategy that allow bacteria to better adapt and survive to changing environments, regulating genes involved in mutagenesis, virulence, antibiotic resistance dissemination, horizontal gene transfer, and inter-species competition [50, 66]. Our work demonstrates that this pathway contributes to *B. cenocepacia* intracellular survival within ciliates and the subsequent enhanced stress resistance and antibiotic tolerance of egested bacteria. Importantly, because the SOS response is a conserved mechanism, our study suggests that protozoan food vacuoles likely constitute an important reservoir that promotes the switch to persister in bacteria that survive intracellular digestion. Furthermore, it has been recently shown that oxidative radicals produced within protozoan food vacuoles facilitate genetic diversification in *V. cholerae* through SOS-induced DNA integration [67]. Exchange of antibiotic resistance genes (ARGs) has been shown to be favored in protozoan food vacuoles, where different bacterial species accumulate under conditions that favor major mechanisms for horizontal gene transfer [67–72]. Therefore, over the last years evidence is accumulating about the role of protozoan food vacuoles as privileged niches that foster genetic diversification and adaptation in bacteria. Deciphering the molecular mechanisms governing this phenomenon is crucial for the development of novel strategies to mitigate pathogen dissemination by modulating bacteria-protist interactions.

## Acknowledgements

This work was supported by a grant from the Spanish Ministry of Science and Innovation, grant number PID2020-113540GB-100 awarded to F.A. A.M. and I.B. were supported by research contracts (PEJ-2021-AI/SAL-21691 and CT36/22-37-UCM-INV, respectively) from the Madrid Government (NextGeneration EU funds) awarded to F.A. We thank Dr. Teresa M. Coque (IRYCIS, Spain) and Natalia Guerra (IRYCIS, Spain) for providing water samples from sink drains, Dr. María Jesús Grilló (IDAB-CSIC, Spain) and Dr. Alicia Sánchez-Gorostiaga (iMiDRA, Spain) for kindly providing strains HB101/pRK2013 and BW2514, respectively. Plasmids pGPI-SceI, pDAI-SceI-SacB and pMLS7eGFP were a gift from Miguel Valvano (Addgene plasmids #32060, #113635 and #112917). Plasmid pSCrhaB2plus was a gift from Silvia Cardona (Addgene plasmid #164226). Plasmid pUC18T-mini-Tn7T-Tp-dsRed was a gift from Herbert Schweizer (Addgene plasmid #65036).

