## Supplementary material for "Protozoan predation enhances stress resistance and antibiotic tolerance in the opportunistic pathogen *Burkholderia cenocepacia* by triggering the SOS response"

Department of Genetics, Physiology and Microbiology. Faculty of Biological Sciences, Complutense University of Madrid. 28040 Madrid. Spain.

**Supplementary Figures**

**Figure S1.** TEM micrographs of wild *Tetrahymena sp* T2305B2 grazing on *B. cenocepacia* K56-2. **(A)** Ciliate expelling EFVs (black arrows) laden with bacteria. **(B)** Aggregate of several EFVs (white arrows) and free bacteria.

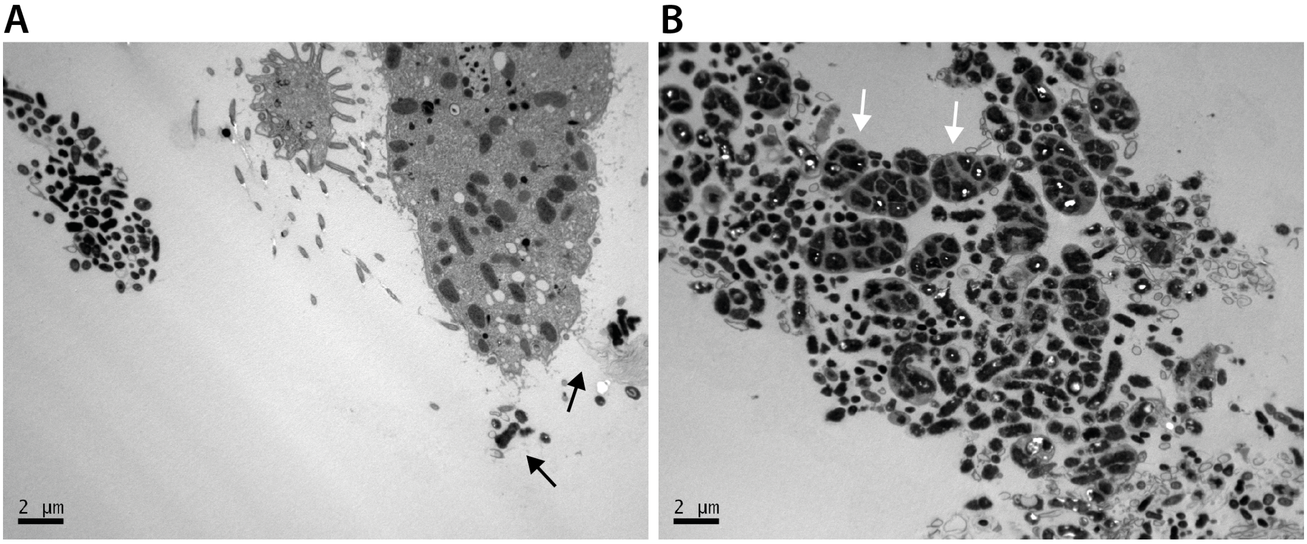

**Figure S2.** Upregulation of SOS response genes in *B. cepacia* ATCC25416 recovered from *T. elliotti* EFVs (24 h post-feeding). Ciliates were incubated with bacteria (ratio 1:200 protist:bacteria) in buffer. At 0 and 24 h ciliates and EFVs were lysed with 1% Triton-X100 and RNA was isolated from collected bacteria. Bacteria incubated in Tris-HCl for 24 h in the absence of ciliates served as control. The stars indicate significant differences (** $p < 0.01$) when compared to the control group (bacteria incubated in buffer for 24 h) according the unpaired t student's test.

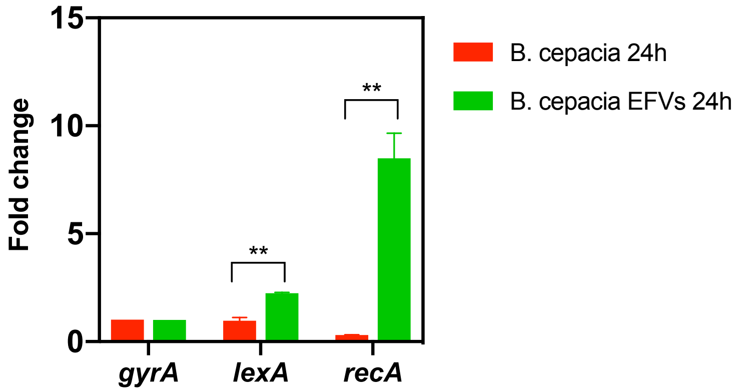

**Figure S3.** Paraquat (PQ) treatment in LB. **(A)** MIC assay for *B. cenocepacia* K56-2 exposed to different concentrations of PQ in LB for 24 h. Bacterial growth was estimated by measuring $OD_{600}$ at 24 h. **(B)** Exposure to 1 mM PQ in LB for 30 min did not result in bacterial death. The number of surviving K56-2 bacteria was quantified by CFU counts on LB agar plates at 0 and 30 min after exposure to PQ.

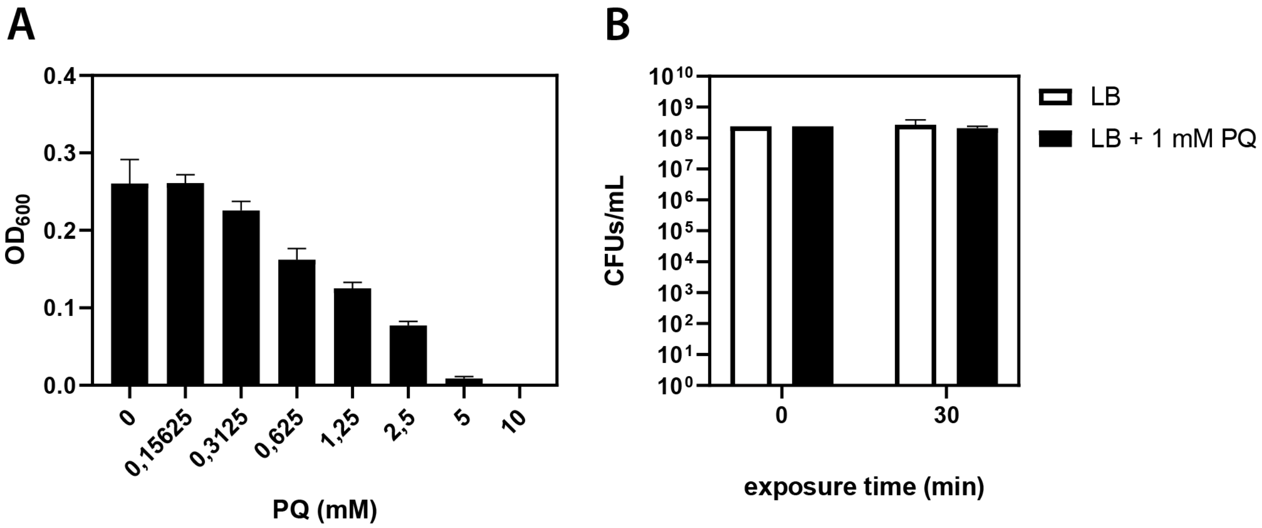

**Figure S4.** Corroboration of the SOS-deficient phenotype of mutant strains Δ*recA* and *lexA(Ind-).* **(A)** Growth curves of strains K56-2, Δ*recA* and *lexA(Ind-)* in LB medium. **(B)** Survival assays of bacteria exposed to UVC (100 $W/cm^2$). SOS-deficient mutants exhibited higher susceptibility to UV compared to parental strain K56-2. Complementation of the *recA* gene in Δ*recA* strain restored UV resistance to wild-type levels. **(C)** Expression levels of *recA* and *lexA* genes in bacteria treated with 2 µg/mL mitomycin C (MMC). MMC failed to induce the SOS response (upregulation of *recA* and *lexA* genes) in strains Δ*recA* and *lexA(Ind-).* Data represent the average of at least three independent biological replicates. Significant differences (* p < 0.05, ** p < 0.01, *** p < 0.005) were calculated using one-way ANOVA and Dunnett's multiple comparison test.

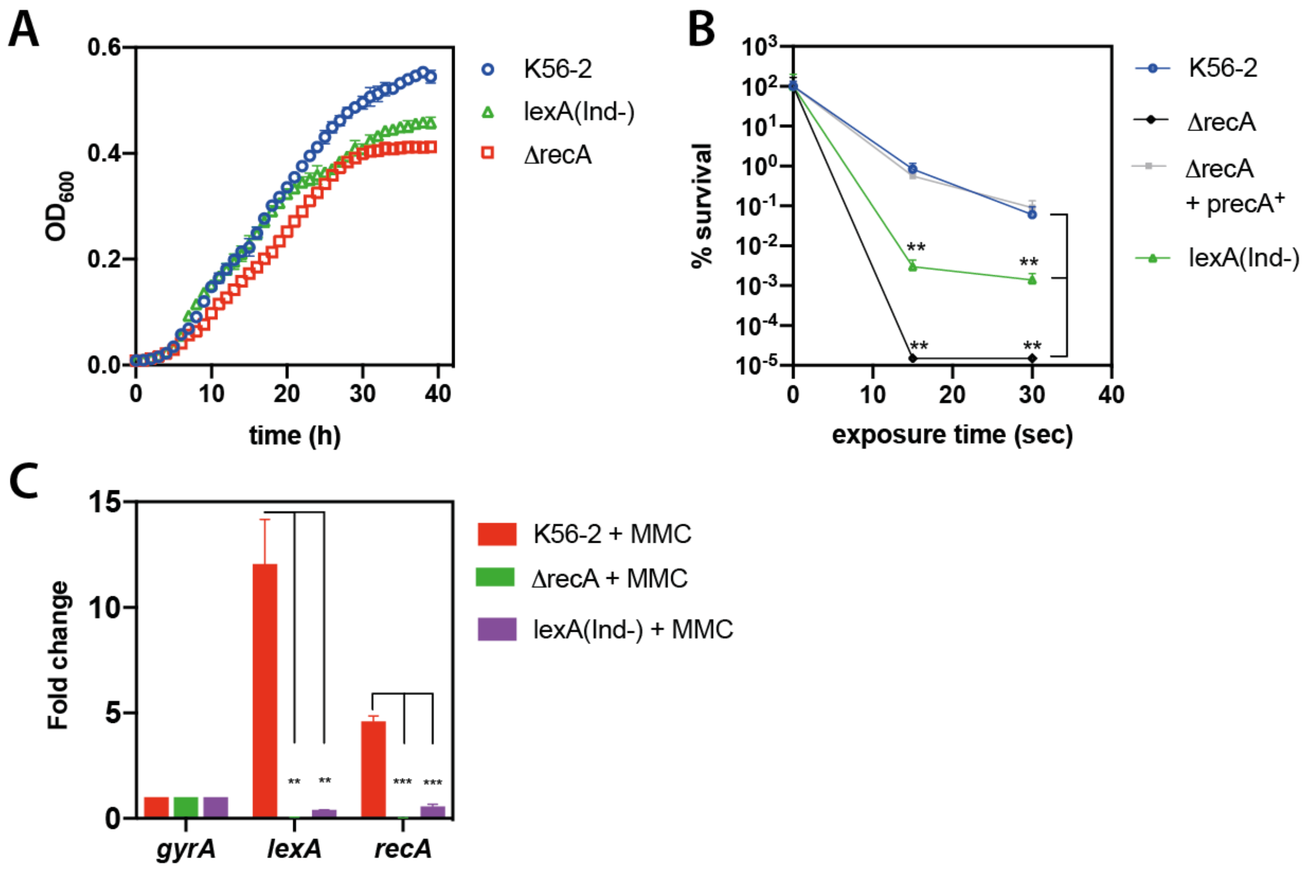

**Figure S5.** Ciprofloxacin minimal inhibitory concentration (MIC) assay of SOS-deficient mutants (Δ*recA* and *lexAInd-*), Δ*surA* and parental strain K56-2. MIC was measured by performing the microdilution test and following CLSI reference methods [1]. Bacterial growth was estimated by measuring $OD_{600}$ at 24 h. MIC was determined as 4 μg/mL for K56-2, 2 μg/mL for *lexA(Ind-)*, <2 μg/mL for Δ*surA* and 1 μg/mL for Δ*recA.*

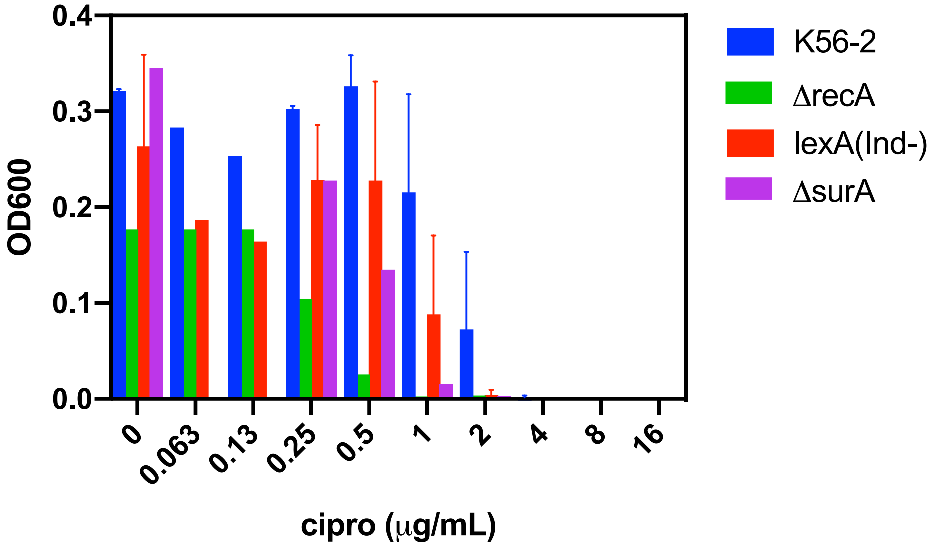

**Table S1.** Bacterial strains and protists used in this study

| Strain | Description | Source |
|---|---|---|
| Bacteria | | |
| *E. coli* | | |
| HB101/pRK2013 | HB101 strain harbouring pRK2013 | M. J. Grillo (IDAB-CSIC) |
| BW25141 | F-, *Δ(araD-araB)567*, *ΔlacZ4787*(*::rrnB*-3), *Δ(phoB-phoR)580*, *λ*$^-$, *galU95*, *ΔuidA3::pir*$^+$, *recA1*, *endA9*(del-ins)*::FRT*, *rph-1*, *Δ(rhaD-rhaB)568*, *hsdR514* | A. Sánchez-Gorostiaga (iMiDRA, Spain) |
| *B. cenocepacia* | | |
| K56-2 | Clinical isolate from a CF patient, ET12 lineage | BCCM/LMG Bacterial Collection |
| Δ*recA* | Δ*recA* derivative of K56-2 | This study |
| FA108 | Δ*recA* harbouring pFA107 plasmid expressing WT *recA* allele under the control of PrhaBAD promoter | This study |
| *lexA*(*Ind-*) | *lexA*(Ind-) derivative of K56-2 | This study |
| FA115 | K56-2 expressing dsRed under the control of PrhaBAD promoter | This study |
| FA161 | K56-2 harbouring pFA160 expressing eGFP under the control of PrecA promoter | This study |
| Δ*surA* | Δ*surA* derivative of K56-2 | [2] |
| *B. cepacia* | | |
| ATCC25416 | Environmental isolate (source *Allium cepa*) | Spanish Type Culture Collection (CECT) |
| Protists | | |
| *Tetrahymena elliotti* 4EA | Ciliate, laboratory strain | Tetrahymena Stock Center Source: environment |
| *Tetrahymena* sp T2305B2 | Ciliate, wild isolate | Hospital sink drain (F. Amaro, unpublished) |
| *Colpoda* sp CE36 | Ciliate, wild isolate | Domestic sink drain (F. Amaro, unpublished) |

**Table S2.** Plasmids used in this study

| Plasmid | Description | Reference |
|---|---|---|
| pRK2013 | $ori_{colE1}$, RK2 derivative, $Kan^r$ $mob^+$ $tra^+$ | [3] |
| pGPI-SceI | $ori_{R6K}$, $mob^+$,$Tp^r$, I-SceI-I restriction site | [4] |
| pDAI-SceI-SacB | $ori_{pBBR1}$, $mob^+$,$Tet^r$, expressing I-Sce-I and sacB | [4] |
| pSCrhaB2plus | pSCrhaB2 with rhaI1-rhaI1 permutation of rhaS binding sites upstream of PrhaBAD and bacteriophage T7 gene 10 stem loop inserted upstream of native rhaBAD 5′ UTR | [5] |
| pFA094 | pGPI-SceI with regions flanking K56-2 *recA* | This study |
| pFA107 | pSCrhaB2plus with K56-2 *recA* gene cloned | This study |
| pFA118 | pGPI-SceI with *lexA*(S119)::$Tet^r$ | This study |
| pFA160 | pPrecAeGFP | This study |

**Table S3.** Oligonucleotides used in this study. Oligonucleotides were synthesized by Integrated DNA Technologies.

| Primer name | Sequence (5' → 3') | Application |
|---|---|---|
| FA198 | TACGTCTAGAAGGATCAGCAGGTCGAAAGT | Generation of plasmids and mutant strains |
| FA199 | TTTTCTCGAGAGCGTGAGCGTGGTCTTA | |
| FA194 | TTTTCTCGAGTGTACGGCGAAGGCATTT | |
| FA200 | TTTTGATATCGCTGCGCGTATAGACATCTT | |
| FA219 | ATTAGACCATATGACCGCCGAGAAGAGCAAG | |
| FA220 | CG TCTAGA TCACTCTTCTTCGTCCATCG | |
| FA223 | GGCTCTGCTGTAGTGAGTGG | |
| FA224 | GCGAAGAAGTTGTCCATATTG | |
| FA225 | TCGCATCTAGACCGTGCGACTGCTTGAAAT | |
| FA226 | GCAACCCACTCACTACAGCAGAGCCTCAGAGTTCGCCCGAGC | |
| FA227 | TGGCCAATATGGACAACTTCTTCGCGTCTCGCTCAGGAGAACAT | |
| FA228 | TTTTGAATTCGACGCGTACGCATAGCC | |
| FA229 | GCGCGGCCTGGCGATGCGCGA | |
| FA230 | ACCTTCAGCAGGTAGTCGGGCTTGCTGG | |
| FA236 | GCGTCGAACCTCAAGTAACA | |
| FA237 | ACGTTGAAGGACCGAGAAAG | |
| FA265 | TTTTCATATGGCCTCCTCCGAGAACGTC | |
| FA266 | TCGATCTAGACTACAGGAACAGGTGGTG | |
| FA289 | TTTATCGATCGAACGGCTTGGCCATGTA | |
| FA290 | TTTCCATGGCCCGGAGCCCTTCTTGCTAT | |
| gyrAFw | CGACGCAGATGAAGGAAGA | Quantitative RT-PCR |
| gyrARv | CATAGACCTTGACCCAGTACAC | |
| lexAFw | GAAGGACGGCCAGATCAT | |
| lexARv | TTTCGTAATCCGGGTTCTCC | |
| recAFw | TGTACGGCGAAGGCATTT | |
| recARv | CGATCTTCTCGCCGTTGTAG | |
| rpoSFw | CGTGATCCGCGAACTGAA | |
| rpoSRv | GGTCTTGCCGGTGAGATAG | |
| oxyRFw | TCGTCGCTCGAAACCATTC | |
| oxyRRv | GCACGTACGACAGCAGTT | |
| katBFw | AGTGGAGCAACGACTTCTTC | |
| katBRv | AATCACCTCGTCCGCATC | |
| sodCFw | TTCCTCGTCGCACGAAA | |
| sodCRv | CGACCAGGTTGTAGGTGAC | |

**Supplementary Methods**

**Determination of MIC.** The MIC was measured by performing the microdilution test and following CLSI reference methods [1]. Bacterial growth was estimated by measuring $OD_{600}$ at 24 h with a TECAN Infinite MPlex plate.

**UV stress assays.** Log-phase cultures ($OD_{600}$=0.3) of wild type and mutant strains were washed and suspended in fresh LB medium. Serial 10-fold dilutions were prepared in fresh LB medium and spotted on the surface of LB agar plates. The plates were then irradiated with 222 nm UVC light at a dose of 100 $W/cm^2$ for different times using a UVO-Ccrosslinker (EQUILAB, SL, Madrid). After irradiation, plates were incubated at 37 ºC for 24 h. The survival percentage of the population was calculated as the number of CFUs divided by the total number of CFUs in the non-irradiated population.
